# The Role of Extended Range of Interactions in the Dynamics of Interacting Molecular Motors

**DOI:** 10.1101/2021.11.09.467943

**Authors:** Cade Spaulding, Hamid Teimouri, S.L. Narasimhan, Anatoly B. Kolomeisky

**Affiliations:** Department of Chemistry, Rice University, Houston, Texas 77005, United States; Center for Theoretical Biological Physics, Rice University, Houston, Texas 77005, United States; Department of Chemical and Biomolecular Engineering, Rice University, Houston, Texas 77005, United States; Chennai Mathematical Institute, SIPCOT IT Park, Siruseri - 603103, India; Department of Physics and Astronomy, Rice University, Houston, Texas 77005, USA

## Abstract

Motor proteins, also known as biological molecular motors, play important roles in various intracellular processes. Experimental investigations suggest that molecular motors interact with each other during the cellular transport, but the nature of such interactions remains not well understood. Stimulated by these observations, we present a theoretical study aimed to understand the effect of the range of interactions on dynamics of interacting molecular motors. For this purpose, we develop a new version of the totally asymmetric simple exclusion processes in which nearest-neighbor as well as the next nearest-neighbor interactions are taken into account in a thermodynamically consistent way. A theoretical framework based on a cluster mean-field approximation, which partially takes correlations into account, is developed to evaluate the stationary properties of the system. It is found that fundamental current-density relations in the system strongly depend on the strength and the sign of interactions, as well as on the range of interactions. For repulsive interactions stronger than some critical value, increasing the range of interactions leads to a change from unimodal to trimodal dependence in the flux-density fundamental diagram. Theoretical calculations are tested with extensive Monte Carlo computer simulations. Although in most ranges of parameters excellent agreement between theoretical predictions and computer simulations is observed, there are situations when the cluster mean-field approach fails to describe properly the dynamics in the system. Theoretical arguments to explain these observations are presented. Our theoretical analysis clarifies the microscopic picture of how the range of interactions influences the dynamics of interacting molecular motors.

## 1. Introduction

It is known that multiple intracellular processes are supported by special classes of active molecules known as motor proteins or biological molecular motors [1,3,6,14]. Such processes include transport along cytoskeleton filaments, cell division, DNA replication and error correction, RNA transcription, synthesis of new protein molecules and many others [1,3,6,13–15]. Molecular motors are frequently viewed as nanomachines that efficiently transform the chemical energy energy into mechanical work needed for their functioning. In recent years, a significant progress in understanding the microscopic mechanisms of individual motor proteins has been achieved from both experimental and theoretical points of view [6,13–15,28]. However, in cells the biological molecular motors typically work in groups, achieving high efficiency by coordinating their activities [14,26]. At the same time, the molecular mechanisms of such cooperativity still remain not well understood [13,20,26,27].

There are several experimental studies suggesting that some motor proteins might interact with each other during the transportation along the protein filaments [23,25,29]. The conclusions from these investigations, however, are rather controversial. Some of them claimed that the interactions for kinesin motor proteins that move along microtubule filaments are weakly attractive [29], while others concluded that these interactions are stronger and more repulsive [25]. But the nature of such interactions and their ranges remain not well explained.

Because most of the motor proteins typically move in a linear fashion, to understand their collective dynamics a powerful method of one-dimensional non-equilibrium multi-particle models, which are known as asymmetric simple exclusion processes, has been widely explored in theoretical studies [5,10,12, 16,20,21]. This approach has been extended to interacting molecular motors that lead to the development of totally asymmetric exclusion process (TASEP) in which interactions are taken into account in a thermodynamically consistent fashion [4,24]. It was later generalized to more realistic situations of biological transport [17–19]. Several other theoretical ideas to analyze the dynamics of interacting particles have been also proposed [2,7–9,11,12,21,22]. This allowed researchers to obtain several new unexpected results concerning the collective dynamics of interacting molecular motors. For example, it was argued that the particle flux through the system might increase for weak repulsive forces in comparison with no interaction case. However, all current theoretical approaches consider only short range interactions between the molecular motors, despite the fact that because of electrostatics (charged groups on motor proteins) such interactions might have longer ranges.

In this paper, we present a theoretical investigation on the role of the range of interactions on dynamics of interacting molecular motors. More specifically, we considered a new TASEP model where not only nearest neighbors but also next-nearest-neighbors interactions are considered. Analysis of this model using a cluster mean-field approach that partially accounts for correlations in the system allows us to obtain a comprehensive description of dynamic properties of the system. Theoretical analysis shows that extending the range of interactions brings a new physics into the system. It is found that a complex trimodal behavior is observed for fundamental current-density diagrams for strong repulsive interactions, while for attractions and weak repulsions a unimodal current-density relation is observed. This is a consequence of the longer range of interactions. We also estimated that the correlations between next-nearest neighbors are weaker and more short-range for repulsive interactions while for attractions they are stronger and more long-range. Theoretical predictions are probed with extensive Monte Carlo computer simulations, and it is found that the theoretical approach correctly describes the dynamics of the system for most ranges of parameters. Several observed deviations are explained due to larger local fluctuations in the particles densities that effectively decrease the correlations in the system. Thus, our analysis suggests that extending the range of interactions introduces qualitatively new phenomena in the complex process of the driven dynamics of interacting particles.

## 2. Theoretical Analysis

### 2.1. Model

Let us consider a model in which transport of molecular motors along linear filaments is viewed as a unidirectional motion of multiple particles on a lattice with *L* (*L* ≫ 1) sites, as shown in Fig. 1 (left). For simplicity, we consider a periodic boundary condition in which site *L* + 1 is the same as the site 1, i.e., the particle dynamics on a ring will be analyzed. Each lattice site *i* (1 ≤ *i* ≤ *L*) is characterized by an occupation number *τ_i_*. If the site *i* is occupied then *τ_i_* = 1, if it is empty then *τ_i_* = 0. Each lattice site can be covered by only one particle.

**Figure 1.**
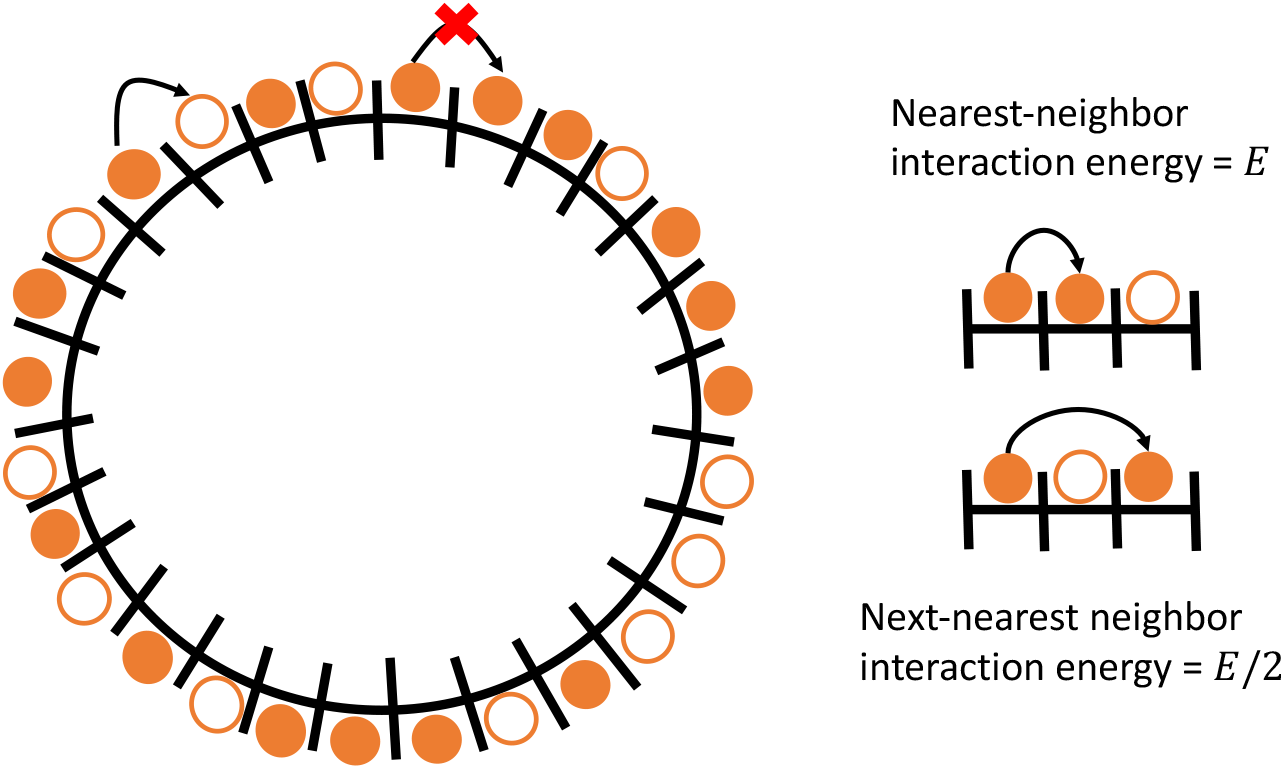
Schematic view of the TASEP model for particles with interactions on a ring of lattice sites (left). Particles move only in the clockwise direction. Filled symbols are particles and empty symbols are holes. Dependence of the interaction energy on the proximity of the neighboring motors (right). The interaction energy for nearest neighbor motors is *E* (in units of *k_B_T*), while the interaction energy for the next-nearest neighbor configuration is *E*/2 (in units of *k_B_T*). See text for more details on the model.

In addition to exclusion, the molecular motors in our model also interact via a long-range potential. Stimulated by the important role of electrostatics, we assume that this potential has the following dependence on the distance *r* between the particles *ϕ*(*r*) ≃ 1/*r*. This means that two motors located *m* sites away from each will have the interaction energy *E/m*, where *E* is the energy of interaction for two nearest neighbors particles (in units of *k_B_T*). It is important to note that *E* < 0 describes the repulsive interactions, while *E* > 0 corresponds to attractions. Previous theoretical studies considered only short-range interactions with *m* = 1. No interactions beyond the nearest neighbors have been considered so far. Our goal is to go beyond *m* = 1, and to clarify the role of the range of interactions on dynamics of interacting molecular motors. For convenience, we concentrate only on the situations that involve nearest neighbors (*m* = 1) and next nearest neighbors (m = 2), neglecting the interactions at longer distances (*m* ≥ 3), although the analysis can be extended to longer ranges of interactions.

The presence of potential modifies the dynamics of creating and breaking ‘bonds’ between the particles. We assume that two particles sitting next to each other form a kind of a ‘stronger bond’, while particles that are separated by one site form a kind of a ‘weaker bond’ due to interactions. The processes of creating and breaking these bonds can be viewed as opposing chemical transitions, which justifies the application of the detailed balance arguments [24]. More specifically, the ratio of forward and backward transition rates between two arbitrary states of the system, labeled *i* and *j*, can be written as

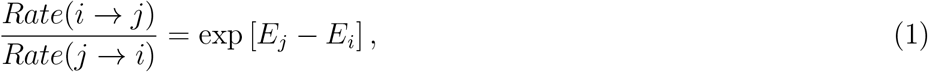

where *E_i_* and *E_j_* are total energies of interactions between all particles in the configurations *i* and *j*, respectively. This means that the ratio of the forward and the backward transition rates between the states *i* and *j* is specified by the difference in energies of these states. To quantify these rates explicitly, we introduce two additional parameters,

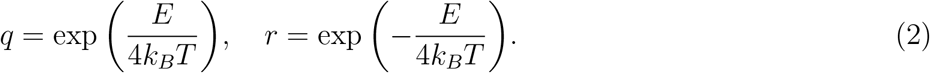

All possible transition rates can be expressed in terms of these two parameters, as we show explicitly below.

### 2.2. Cluster Mean-Field Analysis

It has been argued before that because of interactions there are correlations between particles at neighboring sites in the TASEP model for interacting molecular motors [4,18,19,24]. These correlations are much stronger in comparison with the system without interactions. Neglecting such correlations leads to unphysical behavior for strong interactions between the molecular motors [24]. This indicates that any successful theoretical analysis must take into account these correlations. This is the main reason for us to utilize a theoretical method that is based on *cluster mean-field* calculations.

In this approach, the dynamics inside of the cluster of several lattice sites is considered exactly, while the correlations between different clusters are neglected. Since in our model the interactions do not extend beyond the two lattice sites from the given particle, it seems reasonable to explore a minimal 3-site mean-field cluster analysis. To understand the dynamics in the system, we consider a sequence of six lattice sites, as presented in Fig. 2, where we follow the transitions of the particle from the 3rd site to the 4-th site. Because the lattice ring is homogeneous, this procedure will allow us to evaluate the dynamics at any site of the ring. The stationary probability of a configuration with six sequential sites (*τ_i-2_, τ_i-1_, τ_i_, τ_i+1_, τ_i+2_, τ_i+3_*) is given by *P*(*τ_i-2_, τ_i-1_, τ_i_, τ_i+1_, τ_i+2_, τ_i+3_*). The cluster mean-field approach postulates that the probability of such six-site clusters can be presented as a normalized product of 3-site cluster probabilities [18,19]

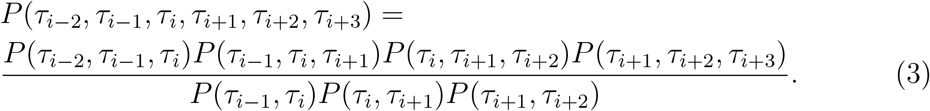

**Figure 2.**
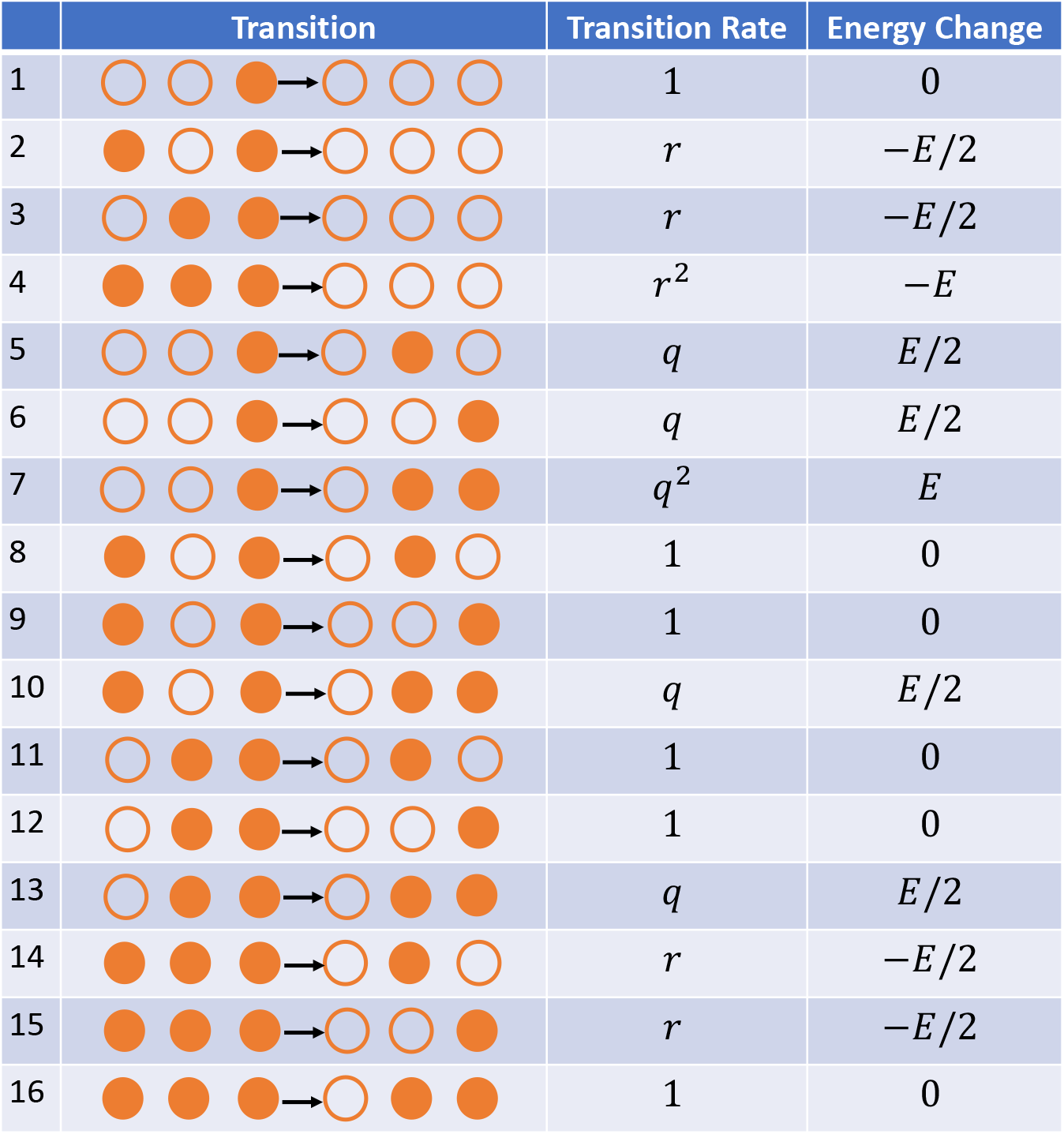
A list of all possible particle transitions, corresponding energy changes and transition rates for a six-site cluster of the lattice. Only the transition of the particle at the third site to the empty fourth is considered.

Note that the denominator accounts for the fact that in the product of 3-site cluster probabilities some sites are accounted twice.

For the six-site cluster there are 16 different transitions as illustrated in Fig. 2. This is because we always start in the configuration with the particle in the 3-rd site and empty 4-th site. Other four sites in the cluster can be in one of two states (occupied or empty), giving the total number of distinguishable configurations to be equal to 2^4^ = 16. Each transition is associated with the energy change and the corresponding transition rates can be found from the detailed balance arguments [see Eq. (1)]. For example, transitions 1, 8, 9, 11, 12 and 16 in Fig. 2 are not associated with any energy changes, and we define the corresponding transition rates to be equal to one. Now let us consider transitions 2 and 6 in Fig. 2. They can be viewed as opposing to each other because in the transition 6 the weak bond is created, while the same weak bond is broken in the transition 2. The energy change associated with this process is just *E*/2. Then the transition rate for the process 2 is set to be equal to r, while for the process 6 it is set to be equal to *q*. This satisfies the detailed balance arguments because *q/r* = exp(E/2). Similar arguments can be applied to all allowed transitions, providing the explicit expressions for different transition rates as shown in Fig. 2.

The overall particle flux in the system will have 16 terms to reflect the different transitions from 16 configurations (see Fig. 2), and it has the following form,

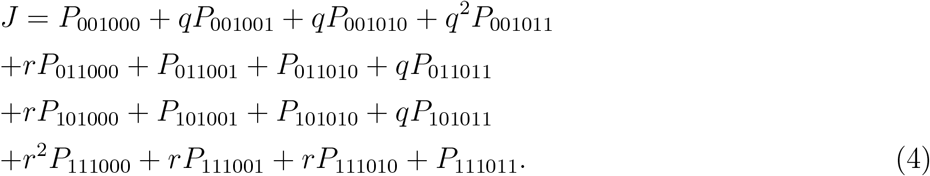

Each probability of the six-site cluster can be expressed via 3-site probabilities using Eq. (3). There are 8 such probabilities labeled as *P*_111_, *P*_110_, *P*_100_, *P*_010_, *P*_011_, *P*_001_ *P*_101_ and *P*_000_. Due to the conservation of probability, they are related via a normalization condition,

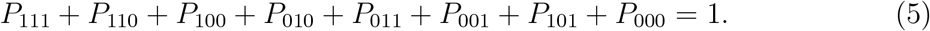

Let us also simplify the notations by defining

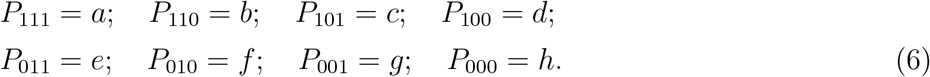

One could also notice that 2-site cluster probabilities can be also expressed via 3-site probabilities,

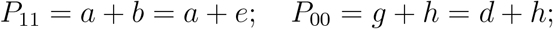

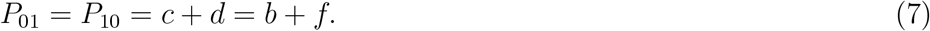

Now, we can obtain an explicit expression for the particle current in the system in terms of the 3-site cluster probabilities,

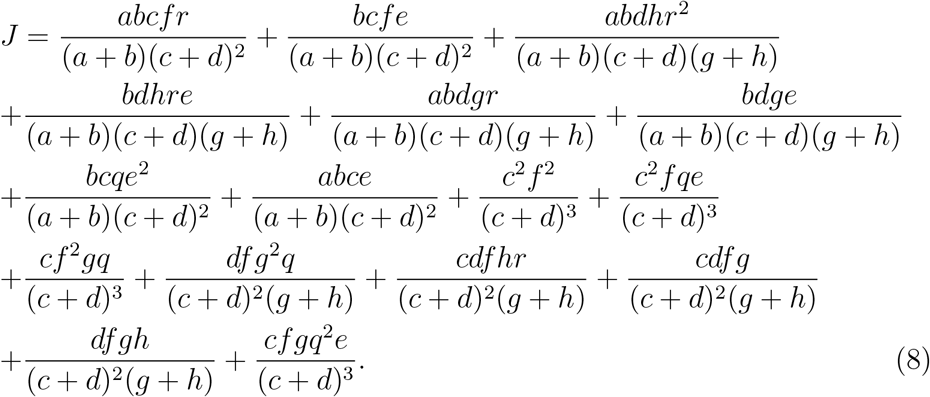

This means that the dynamics in the system of interacting molecular motors can be fully described if we determine all 3-site cluster probabilities.

Thus, we need to explicitly find 8 unknown variables, (*a, b, c, d, e, f, g, h*). To do so, 8 independent equations are required. To find some of these equations, we notice that the system presented in Fig. 1 (left) has a particle-hole symmetry: the clockwise motion of the particles is identical to the counterclockwise motion of the holes. Now, using these symmetry arguments and the normalization, one can easily derive 5 equations:

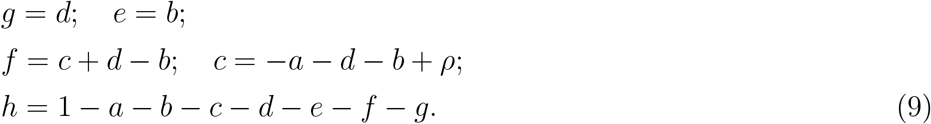

where *ρ* is the bulk density (or the probability to find the particle at a given site). Three more equations are needed, and they can be derived from the corresponding master equations for probabilities *P*_111_, *P*_110_, and *P*_000_,

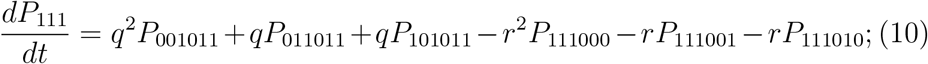

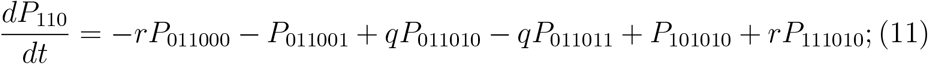

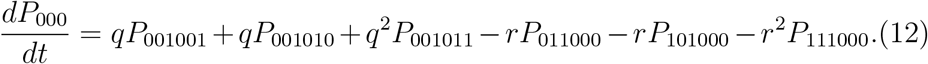

We are interested in the stationary dynamics when the left sides of these equations will be zero, and combining Eqs. (10), (11) and (12) with Eqs. (9) yields a system of three non-linear equations for variables *a*, *b* and *d*,

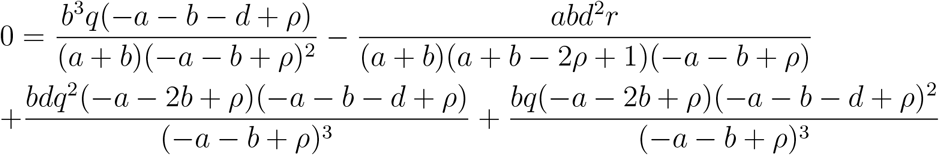

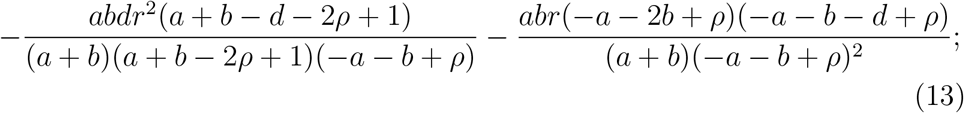

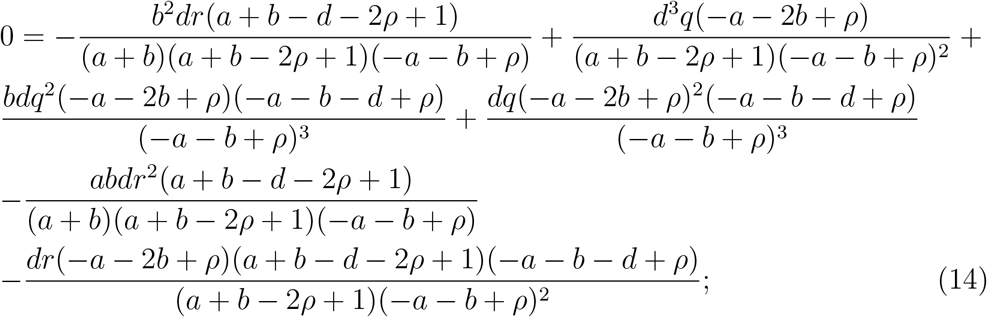

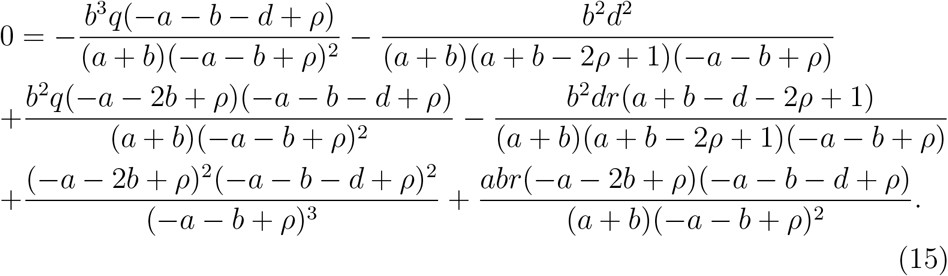

We note that these equations have only two input parameters: the interaction energy *E* and the particle density *ρ*. For any arbitrary value of *E* and *ρ*, Eqs. (9) and (13)-(15) can be solved numerically exactly, providing the explicit expressions for the 3-site cluster probabilities and allowing us to obtain a comprehensive description of the dynamics in the system of interacting molecular motors with extended range of interactions at all possible interactions and particle densities. The results are presented in Fig. 3 where they are also compared with Monte Carlo computer simulations.

**Figure 3.**
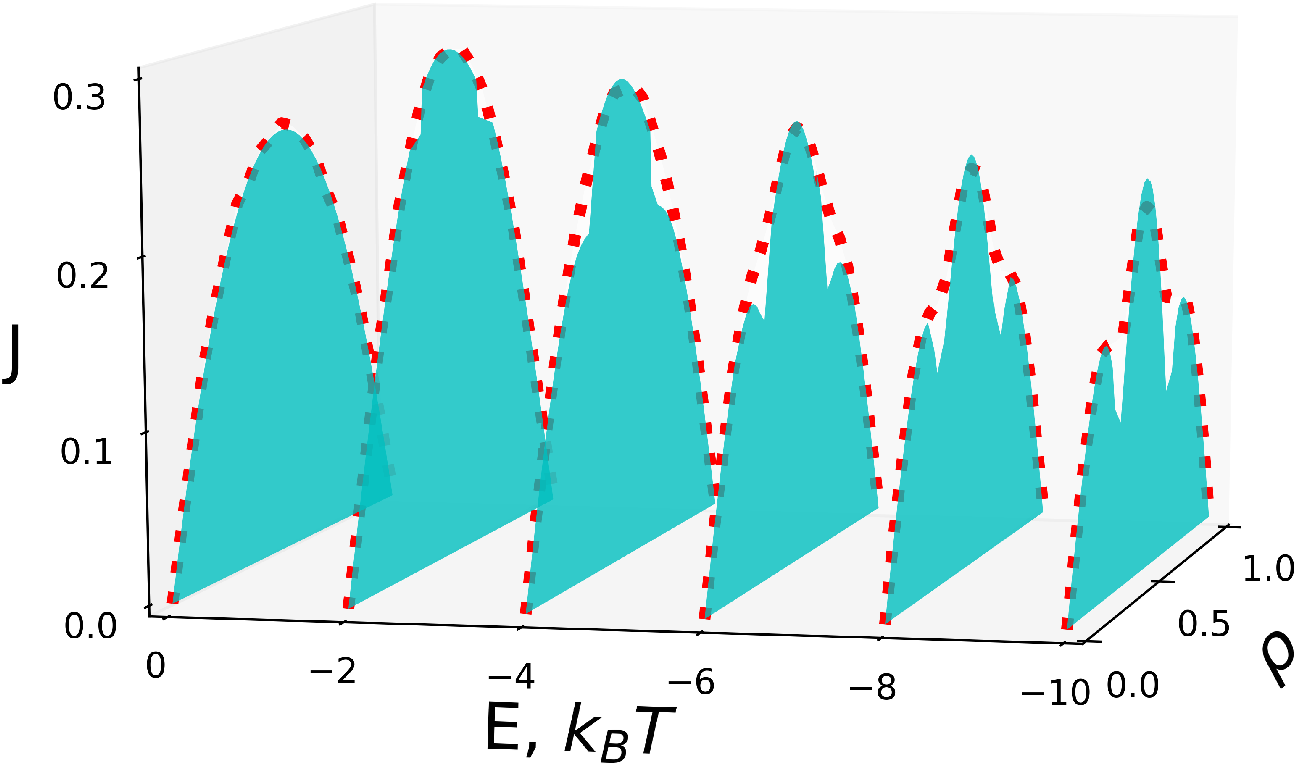
The particle flux in the system of interacting molecular motors for different interactions and densities. Lines are theoretical predictions. Symbols are from Monte Carlo computer simulations. In computer simulations, *L* = 500 was utilized.

### 2.3. Analytical Results for Limiting Situations

Although our theoretical approach can evaluate the dynamics in the system of interacting molecular motors with extended range of interactions numerically exactly, there are several limiting cases when a full analytical description can be obtained. This allows us to prove the validity of our theoretical method as well as to understand better the microscopic picture of these complex processes.

#### 2.3.1. Strongly-Attracting Particles (E → +∞)

In this case, we have *q* → ∞ and *r* → 0, which leads to only two non-zero 3-site cluster probabilities, *P*_111_ = *a* = *ρ* and *P*_000_ = *h* = 1 – *ρ*. Physically it means that all particles on the ring will be jammed in one large continuous cluster that will not allow for any particle flux, *J* = 0. This is an expected behavior because for the particle to break out from the large cluster will cost an infinite amount of energy, and for this reason this does not happen.

#### 2.3.2. Non-interacting Particles (E = 0)

In this limit, we have *q* = *r* = 1, and it can be shown that 3-site cluster probabilities are given by the following expressions:

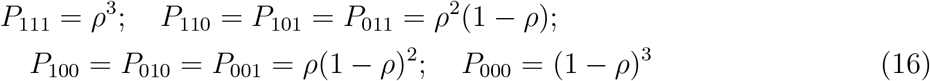

Substituting these results into Eq. (8), we obtain the particle flux as *J* = *ρ*(1 – *ρ*).

These results can be easily interpreted. Without interactions, there is no correlations in the particle occupancy, and the 3-site cluster probabilities can be easily obtained as a product of 3 one-site probabilities with the occupied site probability equal to *ρ* and the empty site probability equal to 1 – *ρ*. These are well-known results for the TASEP model without interactions [5].

#### 2.3.3. Strongly-Repulsive Particles (E → –∞)

In this case, we have *q* → 0 and *r* → ∞, which significantly simplifies the analysis. In addition, it is convenient to perform the calculations for three different ranges of particle densities.

For the first low-density region, when 0 ≤ *ρ* ≤ 1/3, one cannot have 2 particles next to each other due to strong repulsion. Because the density is not high, this can be accomplished, and it leads to

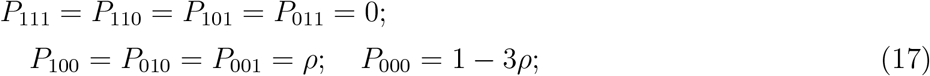

and the following results for the two-site cluster probabilities,

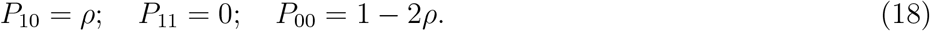

In the limit of strong repulsions, the only contribution to the flux is due to one configuration 001000 out of 16 possible six-site configurations. Thus, the particle flux is given by,

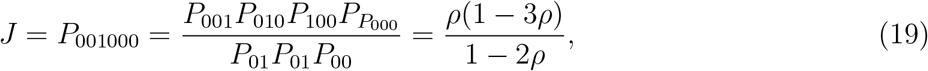

which also corresponds to the flux of non-interacting trimers [17]. Therefore, for 0 ≤ *ρ* ≤ 1/3, for strong repulsions the interacting molecular motors with the increased range of interactions behave like non-interacting timers. This is clearly a result of the increased range that does not allow other particle to come to the given particle closer than 2 sites apart, which effectively mimicks the dynamics of non-interacting trimers.

For intermediate densities, 1/3 ≤ *ρ* ≤ 2/3, it is not possible to avoid the contact of two particles, but the situation with three particles together cannot happen due to the strong repulsions. In this case, we have

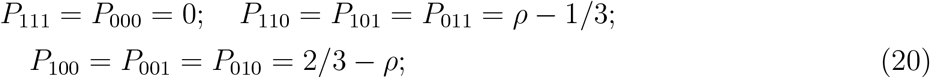

There are four six-site configurations that can produce the particle flux, 101010, 101001, 011010 and 011001. This leads to the following expression for flux,

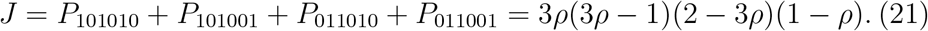

Note that these theoretical arguments predict that the current will disappear for *ρ* = 1/3 and *ρ* = 2/3 in the limit of strong repulsions.

Finally, for the highest density region, 2/3 ≤ *ρ* ≤ 1, the results can be obtained easily using a particle-hole symmetry from the first low-density region calculations. For this purpose, we need to change 0 ↔ 1 and *ρ* ↔ (1 – *ρ*). Then we obtain for the 3-site probabilities,

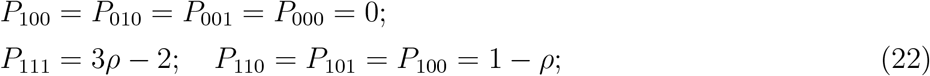

and the following results for the two-site probabilities,

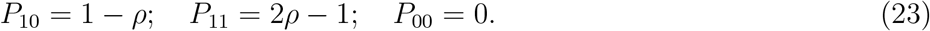

In this case, the only surviving term in particle flux is *P*_111011_. Thus

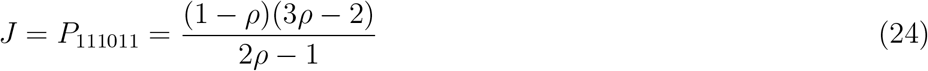

This corresponds to the flux of non-interacting trimers of holes.

Thus, the combined expressions for the particle flux in the strong repulsion limit for interacting molecular motors with extended range of interactions is given by,

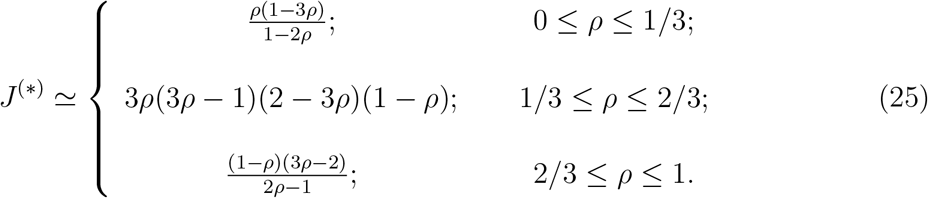

One can see that this function is continuous at 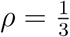 and 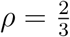, as expected.

### 2.4. Fundamental Current-Density Diagrams

Our theoretical method allows us to explicitly evaluate the relations between the particle currents and densities for all possible inter-molecular interactions, as shown in Fig. 3. It also provides an important insights on the role of the extended range of interactions. A unimodal current-density fundamental diagram is observed for attractions and for weak repulsion with the largest current to be achieved at *ρ* = 1/2. Increasing the strength of repulsion from the no-interactions case *E* = 0 first leads to the increase in the maximal current (at *ρ* = 1/2) at *E* ≃ −2. But the further increase in the repulsion strength leads to the lowering of the maximal current. The interesting behavior starts for repulsions stronger that *E* ≃ −6: the originally unimodal curve with one maximum transforms into the trimodal curve that exhibits three maxima and two minima. The minimum currents are predicted for *ρ* = 1/3 and *ρ* = 2/3. As the repulsion increases, our theory predicts that the currents at these special locations will eventually reach zero, in agreement with our arguments that the system of interacting molecular motors with the extended range of parameters for *E* → –∞ behaves like a system of non-interacting trimers.

We tested the predictions of the 3-site cluster mean-field theory with Monte Carlo computer simulations. Excellent agreement is found for all densities for attractive and weakly repulsive interactions: see Fig. 3. However, there are some deviations from theoretical calculations for strong repulsions near the regions where the minimal current is predicted (around *ρ* = 1/3 and *ρ* = 2/3). Instead of minimal current at these regions, simulations suggest that the particle current becomes almost constant and independent of density, connecting the increasing branches of the current-density curves. This observation will be explained below.

One of the surprising results of the TASEP model with interactions is the observation of the increase in the maximal particle current for weak repulsions [4,24]. But it was found for the system with short-range interactions. Using our theoretical approach, we can probe how the extended range of interactions will affect this result. In Fig. 4, we present the ratio of the particle currents for the system with the extended range (*NNN*) over the particle current for the system with the short-range (*NN*) for different particle densities. For low densities (*ρ* = 0.3) and for high densities (*ρ* = 0.8) there is not much effect on the increasing the flux by extending the range of interactions. However, the strong effect is observed for the maximal current for *ρ* = 1/2. In this case, the flux increases more for the system with extended range of interactions, and the effect is larger for stronger repulsions. One can explain this by noting that for low densities particles can be found at the distances where the interactions do not affect them, while for high densities the particle transitions in most cases are not associated with energy changes. Only for *ρ* =1/2 the effect of interactions is maximal and the extended range helps to make it even stronger.

**Figure 4.**
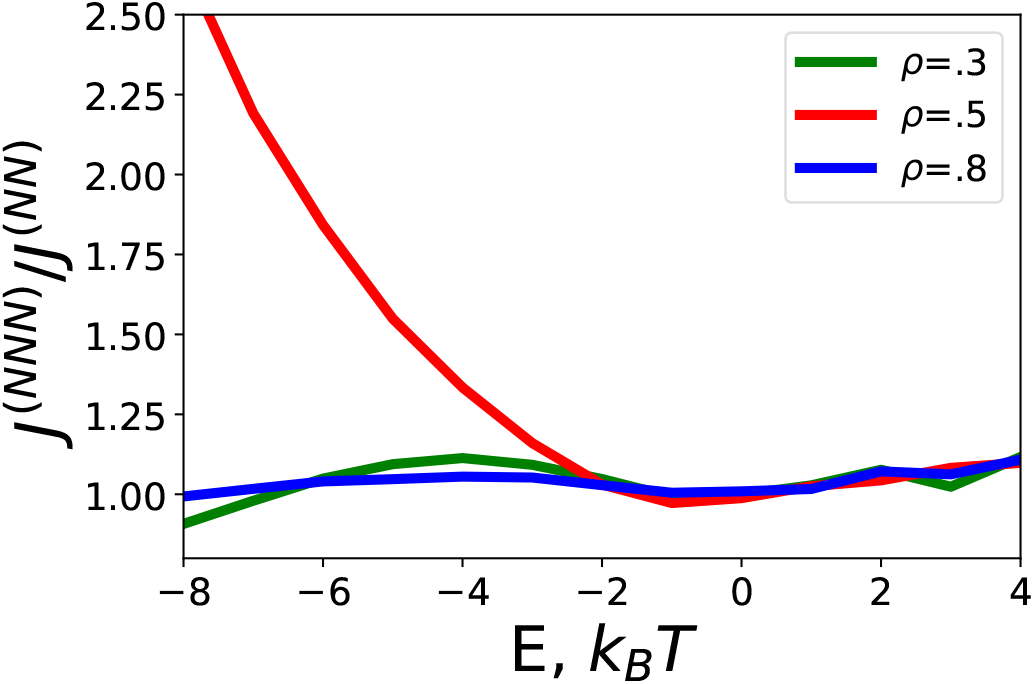
Predictions of Monte-Carlo computer simulations for the ratio of particle fluxes for TASEP with next-nearest neighbor interactions *J^(NNN)^* (extended rage) and TASEP with nearest-neighbor interactions *J^(NN)^* (short range). In simulations, *L* = 500 was utilized.

## 3. Correlation Functions

To understand better the microscopic mechanisms of the processes that are taking place during the transport of interacting molecular motors, it is important to analyze the correlations functions that measure the degree of correlations in the system. One could define the nearest-neighbor correlation function,

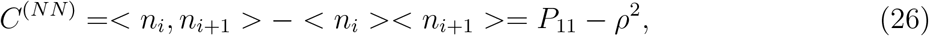

which can be easily calculated using our theoretical approach. However, due to the extended range of interactions in our system, this correlation function might not properly capture the relevant processes in the system. Alternatively, we introduce the next-nearest neighbor correlation function *C^(NNN)^*,

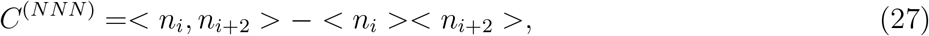

where the corresponding two-point function < *n_i_, n_i+2_* > is defined as

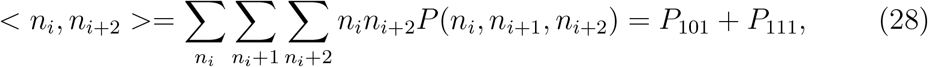

and < *n_i_* >=< *n_i+2_* >= *ρ*. Thus, *C^(NNN)^* is given by

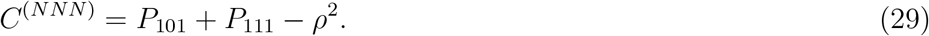

One can see that the main difference between the nearest-neighbor correlation function and the next-nearest neighbor correlation function is that *C^(NNN)^* is expressed in terms of 3-site clusters probabilities while *C^(NN)^* includes only the 2-site clusters probabilities.

Fig. 5 presents the next-nearest neighbor correlation functions, as estimated from the Monte Carlo simulations, as a function of the interactions for different particle densities. One can see that for attractive interactions these correlations are positive, while for repulsions they are negative. These results can be easily explained. For *E* > 0 it is more probable to find another particle at the nearest-neighbor site due to attractive interactions, and this leads to positive correlations. Similarly, for *E* < 0 it is less probable to find the particle at the nearest-neighbor site due to repulsion, and this leads to the negative correlations. It is important to note that these correlations are due to the extended range of interactions and they are essentially absent in the system with short-range interactions.

**Figure 5.**
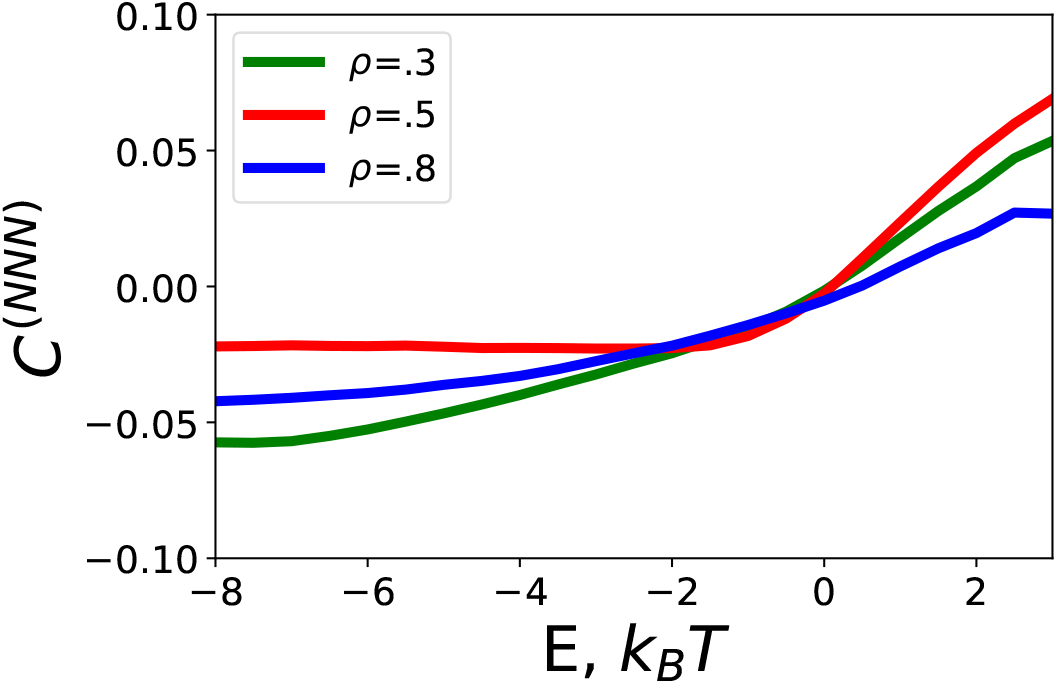
Results of Monte-Carlo simulations for the next-nearest neighbor correlation function *C^(NNN)^* as a function of the interaction energy (*E*). In simulations, *L* = 500 was utilized.

One might illustrate better the effect of the range of interactions by considering several limiting case where these correlation functions can be estimated fully analytically. For the case of no interactions (*E* = 0), it was already shown that *P*_111_ = *ρ*^3^ and *P*_111_ = *ρ*^2^(1 – *ρ*). The substituting these results into Eq. (29) produces *C^(NNN)^* = 0, i.e., there are no next-nearest neighbor interactions, as expected. Analytical results can be also obtained in the limit of strong repulsions, *E* → –∞. Using Eq. (17), we derive

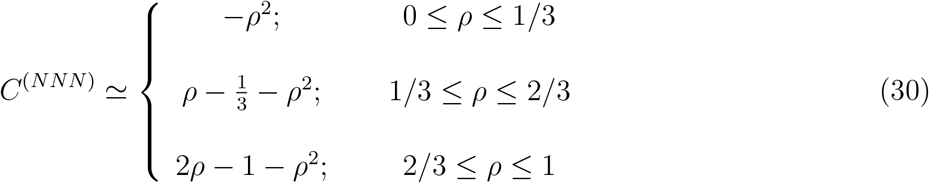

Our theoretical 3-site cluster mean-field approach is successful in describing the dynamics of interacting molecular motors with the extended range of interactions for attractions and weak repulsions. Only for strong repulsions, the deviations between computer simulations and theoretical predictions are found near the regions of *ρ* = 1/3 and *ρ* = 2/3. In these cases, theory predicts a minimal particle current which is not observed in Monte Carlo simulations at all. To understand the origin of such deviations, we plot in Fig. 6 the particle flux and the next-nearest neighbor correlation function for very strong repulsion (*E* = –20). The Monte Carlo computer simulations are comparedwith the calculations of the 3-site cluster mean-field approach for interacting particles with the extended range and for non-interacting trimers [18]. This is because for such strong repulsions one could expect that the dynamics of non-interacting trimers might provide a good description, as we argued above.

**Figure 6.**
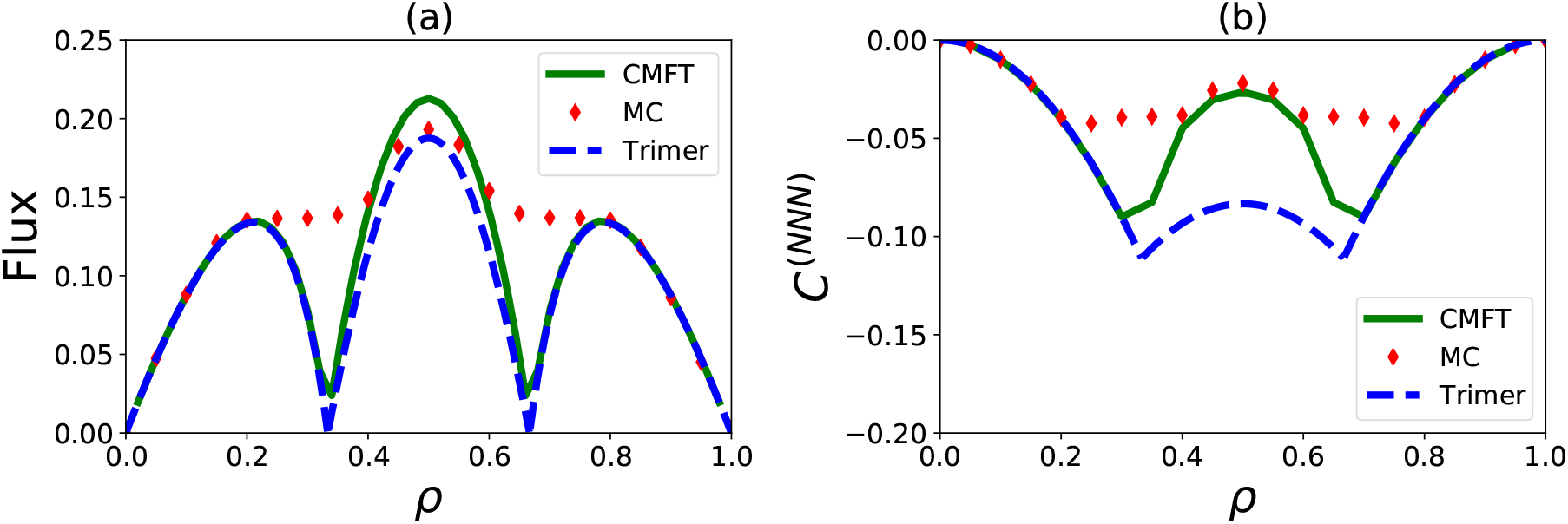
Predictions of Monte-Carlo computer simulations and cluster mean-field theory for *E* = –20*k_B_T* for interacting molecular motors and for non-interacting trimers. (a) The particle flux (*J*) as a function of particle density (*ρ*); and (b) The next-nearest neighbor correlation function (*C^(NNN)^*) versus particle density.

Note that our calculations of the “trimers” theory are done in the following way. For small densities (*ρ* < 1/3) it is assumed that the trimer corresponds to 111 clusters, for intermediate densities (1/3 < *ρ* < 2/3) the trimer describes 110 clusters, while for high densities (2/3 < *ρ* < 1) the trimer corresponds to 100 clusters. We calculate the fluxes of such particles in all different regimes. Of course, in our real system particle follow slightly different dynamics since the monomer transitions are taking place. However, the trimer approach might be a very reasonable description for very strong repulsions.

Comparison of Fig. 6b and Fig. 6a suggests that the theoretical calculations do not work well in the region of minimal currents due to overestimating the negative correlation in the system. In reality, the negative next-nearest neighbor interactions are not as strong. The possible reason for such observations is the following. Let us consider for simplicity the case *ρ* = 1/3. The particle density in the system constantly fluctuates. This means that there are segments where the density is larger than 1/3 and there are segments where the density is smaller than 1/3. But for such densities, high currents will be realized (see branches in the fundamental diagram in Fig. 6a). As the result, the fluctuations will wash out the negative correlations and larger particle fluxes are observed. The cluster mean-field theory does not account for such fluctuations, and this is the reason for differences between theoretical calculations and computer simulations. Note that when the theory correctly captures the correlations (e.g., near *ρ* = 1/2) the agreement between the simulations and theoretical predictions is much better.

## 4. Summary and Conclusions

We presented a theoretical study on the role of the extended range of interactions in dynamics of interacting molecular motors. Our analysis is based on investigating the TASEP model of particles with nearest and next-nearest interactions that move unidirectionally along the lattice of sites. To account for correlations in the system that are driven by interactions we developed a cluster mean-field theory that allowed to evaluate numerically exactly the dynamic properties in the system. Theoretical calculations are also supplemented by extensive Monte Carlo computer simulations. It is found from theoretical calculations that a unimodal fundamental current-density diagrams will be observed for attractive and weak repulsions. For stronger than some critical value of the repulsion, it is predicted that the flux-density relation will exhibit a trimodal shape. This is the result of the extended range of interactions. Computer simulations agree with theoretical predictions for most situations except for the regions near the minimal current for strong repulsions. It is argued that the differences between the theoretical predictions and computer simulations are due to fluctuations that lower the amplitude of correlations at these regions. Our theoretical analysis shows how the extended range of interactions influences the topology of fundamental diagrams, the correlations and the particle currents, clarifying important microscopic features of complex non-equilibrium dynamic processes.

While our theoretical approach considered only the situation when interactions affect only nearest and next-nearest particles, it can be extended for longer ranges of interactions. Based on our analysis presented in this work, we expect that longer ranges of interactions will affect the dynamics only at repulsions. It will lead to the increased maximal current, increased correlations and more complex fundamental diagrams that might exhibit multi-modal shapes that probably will be washed out by fluctuations in the system. It will be interesting to test these predictions in more quantitative theoretical models and more advanced computer simulations.

## Acknowledgments

The work was supported by the Welch Foundation (C-1559), by the NSF (CHE-1953453 and MCB-1941106), and by the Center for Theoretical Biological Physics sponsored by the NSF (PHY-2019745).

## References

[1] Bruce Alberts, Alexander Johnson, Julian Lewis, Martin Raff, Keith Roberts, and Peter Walter. Molecular biology of the cell. 2007.

[2] Tibor Antal and GM Schutz. Asymmetric exclusion process with next-nearest-neighbor interaction: Some comments on traffic flow and a nonequilibrium reentrance transition. Physical Review E, 62(1):83, 2000.

[3] Paul C Bressloff and Jay M Newby. Stochastic models of intracellular transport. Reviews of Modern Physics, 85(1):135, 2013.

[4] Daniel Celis-Garza, Hamid Teimouri, and Anatoly B Kolomeisky. Correlations and symmetry of interactions influence collective dynamics of molecular motors. Journal of Statistical Mechanics: Theory and Experiment, 2015(4):P04013, 2015.

[5] T Chou, K Mallick, and RKP Zia. Non-equilibrium statistical mechanics: from a paradigmatic model to biological transport. Reports on progress in physics, 74(11):116601, 201l.

[6] Debashish Chowdhury. Stochastic mechano-chemical kinetics of molecular motors: a multidisciplinary enterprise from a physicist’s perspective. Physics Reports, 529(1):1–197, 2013.

[7] Marcel Dierl, Mario Einax, and Philipp Maass. One-dimensional transport of interacting particles: Currents, density profiles, phase diagrams, and symmetries. Physical review E, 87(6):062126, 2013.

[8] Marcel Dierl, Philipp Maass, and Mario Einax. Time-dependent density functional theory for driven lattice gas systems with interactions. EPL (Europhysics Letters), 93(5):50003, 2011.

[9] Marcel Dierl, Philipp Maass, and Mario Einax. Classical driven transport in open systems with particle interactions and general couplings to reservoirs. Physical review letters, 108(6):060603, 2012.

[10] Jiajia Dong, Stefan Klumpp, and Royce KP Zia. Entrainment and unit velocity: Surprises in an accelerated exclusion process. Physical review letters, 109(13):130602, 2012.

[11] JS Hager, J Krug, V Popkov, and GM Schutz. Minimal current phase and universal boundary layers in driven diffusive systems. Physical Review E, 63(5):056110, 200l.

[12] Stefan Klumpp and Reinhard Lipowsky. Phase transitions in systems with two species of molecular motors. EPL (Europhysics Letters), 66(1):90, 2004.

[13] Anatoly B Kolomeisky. Motor proteins and molecular motors: how to operate machines at the nanoscale. Journal of Physics: Condensed Matter, 25(46):463101, 2013.

[14] Anatoly B Kolomeisky. Motor proteins and molecular motors. CRC press, 2015.

[15] Anatoly B Kolomeisky and Michael E Fisher. Molecular motors: a theorist’s perspective. Annu. Rev. Phys. Chem., 58:675–695, 2007.

[16] Reinhard Lipowsky, Stefan Klumpp, and Theo M Nieuwenhuizen. Random walks of cytoskeletal motors in open and closed compartments. Physical Review Letters, 87(10):108101, 200l.

[17] Tripti Midha, Luiza VF Gomes, Anatoly B Kolomeisky, and Arvind Kumar Gupta. Theoretical investigations of asymmetric simple exclusion processes for interacting oligomers. Journal of Statistical Mechanics: Theory and Experiment, 2018(5):053209, 2018.

[18] Tripti Midha, Anatoly B Kolomeisky, and Arvind Kumar Gupta. Effect of interactions for one-dimensional asymmetric exclusion processes under periodic and bath-adapted coupling environment. Journal of Statistical Mechanics: Theory and Experiment, 2018(4):043205, 2018.

[19] Tripti Midha, Anatoly B Kolomeisky, and Arvind Kumar Gupta. Interactions in nonconserving driven diffusive systems. Physical Review E, 98(4):042119, 2018.

[20] Izaak Neri, Norbert Kern, and Andrea Parmeggiani. Exclusion processes on networks as models for cytoskeletal transport. New Journal of Physics, 15(8):085005, 2013.

[21] Itai Pinkoviezky and Nir S Gov. Modelling interacting molecular motors with an internal degree of freedom. New Journal of Physics, 15(2):025009, 2013.

[22] Vladislav Popkov and Gunter M Schutz. Steady-state selection in driven diffusive systems with open boundaries. EPL (Europhysics Letters), 48(3):257, 1999.

[23] Wouter H Roos, Otger Campas, Fabien Montel, Gunther Woehlke, Joachim P Spatz, Patricia Bassereau, and Giovanni Cappello. Dynamic kinesin-1 clustering on microtubules due to mutually attractive interactions. Physical biology, 5(4):046004, 2008.

[24] Hamid Teimouri, Anatoly B Kolomeisky, and Kareem Mehrabiani. Theoretical analysis of dynamic processes for interacting molecular motors. Journal of Physics A: Mathematical and Theoretical, 48(6):065001, 2015.

[25] Ivo A Telley, Peter Bieling, and Thomas Surrey. Obstacles on the microtubule reduce the processivity of kinesin-1 in a minimal in vitro system and in cell extract. Biophysical journal, 96(8):3341–3353, 2009.

[26] R TyleráMcLaughlin et al. Collective dynamics of processive cytoskeletal motors. Soft matter, 12(1):14–21, 2016.

[27] Karthik Uppulury, Artem K Efremov, Jonathan W Driver, D Kenneth Jamison, Michael R Diehl, and Anatoly B Kolomeisky. How the interplay between mechanical and nonmechanical interactions affects multiple kinesin dynamics. The Journal of Physical Chemistry B, 116(30):8846–8855, 2012.

[28] Claudia Veigel and Christoph F Schmidt. Moving into the cell: single-molecule studies of molecular motors in complex environments. Nature Reviews Molecular Cell Biology, 12(3):163–176, 2011.

[29] Andrej Vilfan, Erwin Frey, Franz Schwabl, Manfred Thormählen, Young-Hwa Song, and Eckhard Mandelkow. Dynamics and cooperativity of microtubule decoration by the motor protein kinesin. Journal of molecular biology, 312(5):1011–1026, 2001.

